# Bombs and cocaine: detecting nefarious nitrogen sources using remote sensing and machine learning

**DOI:** 10.1101/842641

**Authors:** Christopher Adams, Soo Mei Chee, David Bell, Oliver P.F. Windram

## Abstract

Plants are treated with synthetic or organic nitrogen sources to increase growth and yield, the most common being calcium ammonium nitrate. However, some nitrogen sources are used in illicit activities. Ammonium nitrate is used in explosive manufacture and ammonium sulphate in the cultivation and extraction of the narcotic cocaine from *Erythroxylum spp*. Here we show that hyperspectral sensing, multispectral imaging and machine learning image analysis can be used to visualise and differentiate plants exposed to different nefarious nitrogen sources. Metabolomic analysis of leaves from plants exposed to different nitrogen sources reveals shifts in colourful metabolites that may contribute to altered reflectance signatures. Overall this suggests that different nitrogen feeding regimes alter plant secondary metabolism leading to changes in the reflectance spectrum detectable via machine learning of multispectral data but not the naked eye. Our results could facilitate the development of technologies to monitor illegal activities involving various nitrogen sources and further inform nitrogen application requirements in agriculture.

## Introduction

Synthetic or organic nitrogen sources applied to soils or plant tissue provide essential nutrients to enhance and facilitate plant growth. Variable nitrogen abundance can alter gene expression and metabolic pathways (Stitt, 1999). These pathways alter root growth, influencing nutrient uptake, the structure of the plant and nitrogen distribution in the surrounding environment. Assimilation of nitrogen within plant tissue can vary dependent on the presence of certain enzymes (Foyer, 2018), which in turn are dependent on the nutrient status of the plant. Nitrogen deficiency can shift secondary metabolism from nitrogen rich alkaloids to carbon rich molecules. Overall, the transcriptome, proteome and the metabolome are all influenced by nitrogen availability (Fritz *et al*., 2006).

64% of the 0.86 million tonnes of nitrogen used annually in the UK is in the form of Ammonium Nitrate (AN) or Calcium Ammonium Nitrate (CAN) (Harty *et al*., 2016). Recently, there has been a noticeable shift towards using CAN in agriculture as AN is a detonatable oxidiser that can be used to construct simple yet highly destructive explosives (Brust *et al*., 2015; Djerdjev *et al*., 2018). Infamously, used by terrorists AN is inexpensive^6^ and small amounts are needed to create explosives (Figuli *et al*., 2016), which have been used in numerous attacks (Anil Ananthaswamy, 2004; Henry Dodd, 2018). Subsequently, Ireland has banned the sale of AN based fertilisers (The Government Inspector of Explosives, Ireland, 2016), while the European commission prohibits the distribution of fertilisers containing more than 16% AN (ECHA, 2009). The USA is also introducing new regulation to limit its use (CISA, 2019).

Ammonium sulphate (AS) is another nitrogen source associated with illegal activity. AS is used in the production of cocaine from *Erythroxylum coca* and *Erythroxylum novogranatense* (U.S. Department of Justice, 1991; Mallette *et al*., 2017). *Erythroxylum* species prefer moderately acidic soil conditions (Johnson, Campbell and Foy, 1997), therefore, AS is used lower soil pH enhancing yields (Johnson, Campbell and Foy, 1997). Additionally, the extraction of cocaine from coca leaves produces AS as a by-product after Sulfuric acid is added to coca leaf paste and then washed with Ammonia (U.S. Department of Justice, 1991; Mallette *et al*., 2017).

Remote sensing represents an appealing strategy to monitor dangerous objects or illicit activity as it is non-invasive and has the potential to scale up using airborne sensors. Remote sensing measures shifts in electromagnetic reflectance, absorption and/or fluorescence to infer the chemical nature of objects under study (Schott, 2007). Close range remote sensing has been used to detect explosives such as TNT and DNT (Keith J. Albert *et al*., 2001; Jason M. Conder *et al*., 2003). Similar approaches have also been used to detect cocaine in acetonitrile solution (Wren *et al*., 2014). Furthermore, high resolution spectroscopy has also been developed to detect a wide range of explosives (De Lucia *et al*., 2003; Federici *et al*., 2005; Liu *et al*., 2006) and differentiate cocaine concentrations in caffeine, glucose and rum (Ryder, O’Connor and Glynn, 2000; Eliasson, Macleod and Matousek, 2008). However, these methods require expensive fine-tuned instruments and access to physical samples with their adaption as airborne imaging systems appearing limited.

Determination of nitrogen uptake by plants and nitrogen content in soil is an area of intense interest in agricultural research. Several remote sensing and imaging methods have been developed to estimate the nitrogen status of plants (W. C. Bausch and H. R. Duke, 1996; Serrano, Filella and Pen~uelas, 2000; Smith *et al*., 2003; Schlemmer *et al*., 2005; Pagola *et al*., 2009). However, in general the focus of this research has been on estimating total nitrogen content *in planta* and relating its impact to yield, rather than detection of different nitrogen compound applications. For this satellite and airborne imaging systems have been developed to help estimate nitrogen abundance in crops (Swain, Jayasuriya and Salokhe, 2007; Rama Rao *et al*., 2008).

Given the role of different nitrogen compounds in illicit explosive and narcotic manufacture and that they are readily bioassimulated by plants, we were curious what impact their presence has on plant physiology, and if this could be used to detect these substances. In this paper, we show that different nitrogen sources alter plant leaf reflectance and plant metabolism. Through nitrogen application experiments using *Arabidopsis thaliana, Solanum lycopersicum* (tomato) and mixed *Panicum* species (lawn grass) we reveal with hyperspectral sensing how different nitrogen source applications alter plant leaf reflectance. Then we show how customised multispectral imaging using conventional RGB cameras and machine learning image analysis can distinguish plants exposed to different nitrogen compounds. Finally, we show through metabolomic profiling that different nitrogen compounds alter the plant leaf metabolome and suggest how some of these secondary metabolites might influence plant leaf reflectance.

## Results and Discussion

### Hyperspectral reflectance reveals significant differences in plants exposed to different nefarious nitrogen compounds

To determine if plant leaf reflectance can be used to distinguish between plants exposed to different nitrogen compounds, we first looked at total plant leaf reflectance using hyperspectrometry. Arabidopsis, tomato and lawn grass were treated with equimolar (4mM) amounts of either AN (explosives), AS (cocaine manufacture), CAN (safe agricultural fertiliser) or a water control. One-week later leaf hyperspectral reflectance data was collected for all species (Fig. 1a and Supplementary Fig. 1). As part of the aim of our study was to adapt conventional RGB cameras to facilitate distinction of nitrogen treatments we chose to focus on the visual part of the electromagnetic spectrum (VIS-EM) 400 – 700nm. Reflectance intensity in this region follows a trend across species with relative intensity differences between treatments being hard to distinguish.

**Figure 1.**
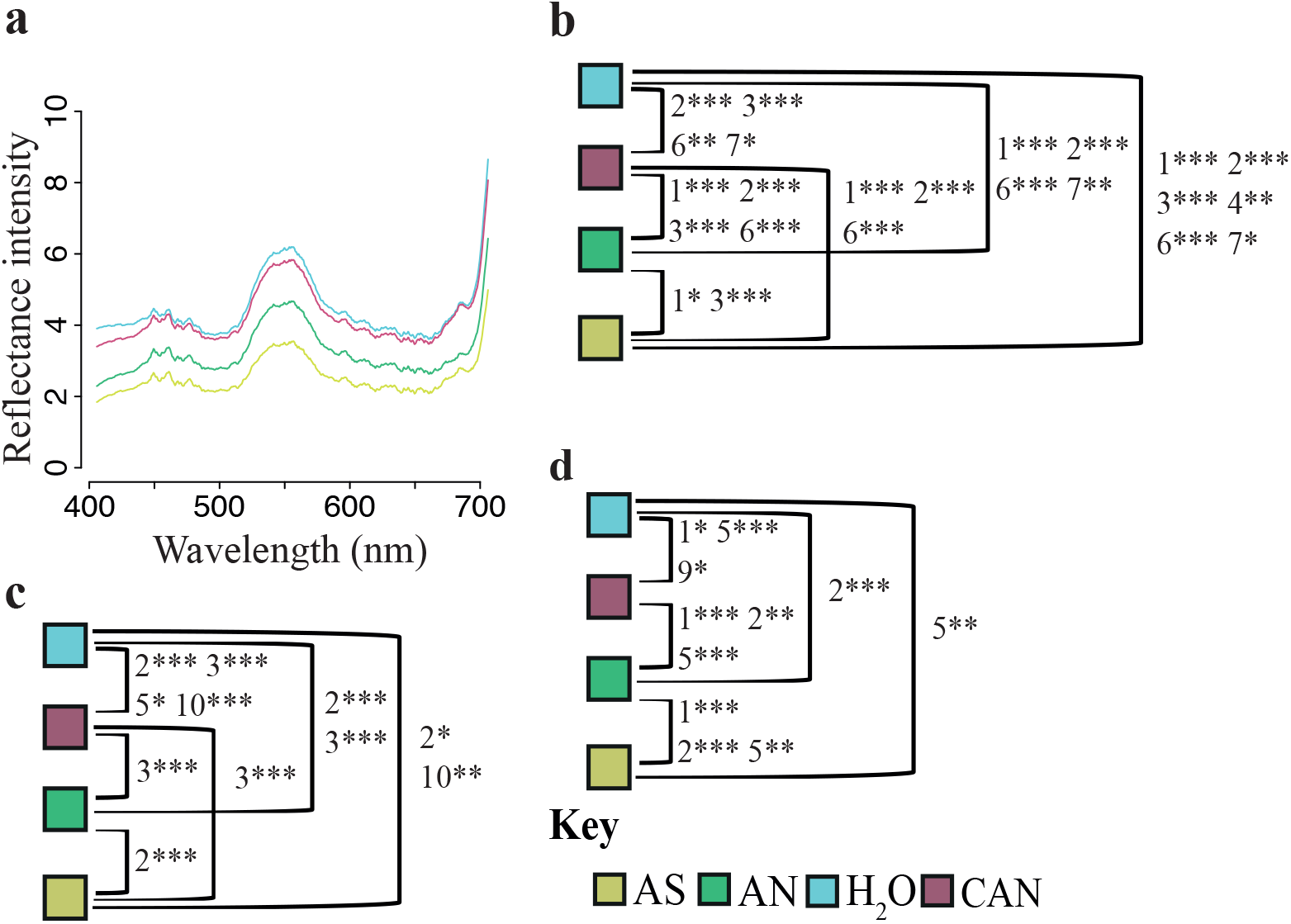
Hyperspectral variance reveals differences in plants treated with different nitrogen sources. Mean reflectance intensity from 400-700nm for **a** tomato one week after application of either 4mM of AN (green), CAN (red), AS (yellow) or H_2_O (blue), 1000 readings taken from 24 plants per treatment. ANOVA and Tukey’s HSD following PCA of reflectance data (400-700 nm) performed on PCs 1-10 **b** tomato **c** Arabidopsis **d** lawn grass. Lines between groups are labelled with PCs which contain variation that can significantly distinguish treatment group pairs. * indicates p-value for Tukey’s HSD < 0.05, ** < 0.01 and *** < 0.001.

We performed principle component analysis (PCA) on VIS-EM hyperspectral reflectance data (Fig 1b, c & d). Overall, in Arabidopsis 93.15%, in tomato-95.74% and in grass-79.23% of the variation can be explained by the first and second principle components (PCs), while in Arabidopsis-94.58%, in tomato-97.53% and in grass-83.79% of total variation is captured in the first 10 PCs (Supplementary fig 2). It is likely that the lower total level of variance explained for grass is influenced by the small size of grass leaves and the need to sample over swards versus single leaves captured for Arabidopsis and tomato.

Next, we analysed the first 10 PCs for each plant by analysis of variance followed by a Tukey’s Honest Significance Difference test (Fig. 1b-d and Supplementary Fig. 3). The four nitrogen treatments can be significantly distinguished by variation captured within the first 10 PCs. It is common practice to perform analysis of variance on the first or second PC often discarding PCs with lower eigenvalues (Climaco Pinto *et al*., 2008; Zwanenburg *et al*., 2011). However, it has been rightly noted (Jolliffe, 1982) that under certain conditions PCs that capture only small amounts of variance can be important. Here we argue that consideration of PCs with small eigenvalues can be important, when distinguishing plant leaf reflectance groups. This is shown in our data where we see PCs with small eigenvalues capable of distinguishing our treatment classes (Fig 1 and Supplementary Fig 2 & 3).

### Ensemble machine learning regression highlights VIS-EM regions with high variability

Having established that sufficient reflectance variation exists within the VIS-EM to distinguish plant leaves treated with different nitrogen compounds, we now wanted to identify key regions to target with multispectral imaging. For this we used an ensemble machine learning approach (Feilhauer, Asner and Martin, 2015), combining partial least square, random forest and support vector machine regression to identify VIS-EM regions exhibiting high spectral variance across nitrogen treatments (Fig. 2). The approach was adapted to identify variance within and across species that differed significantly between nitrogen treatments. This identified multiple VIS-EM regions. In particular, we note that 400-410, 445-455, 540-560, 610 and 675-700 nm are consistently identified across all species. Overall, this highlights a complexity of VIS-EM regions exhibiting altered reflectance patterns across nitrogen treatments, both unique and conserved across species. Supporting this we observe eigenvalues of PC highlighted in our ANOVA derive high variance contribution from these regions (Supplemental Fig. 4).

**Figure 2.**
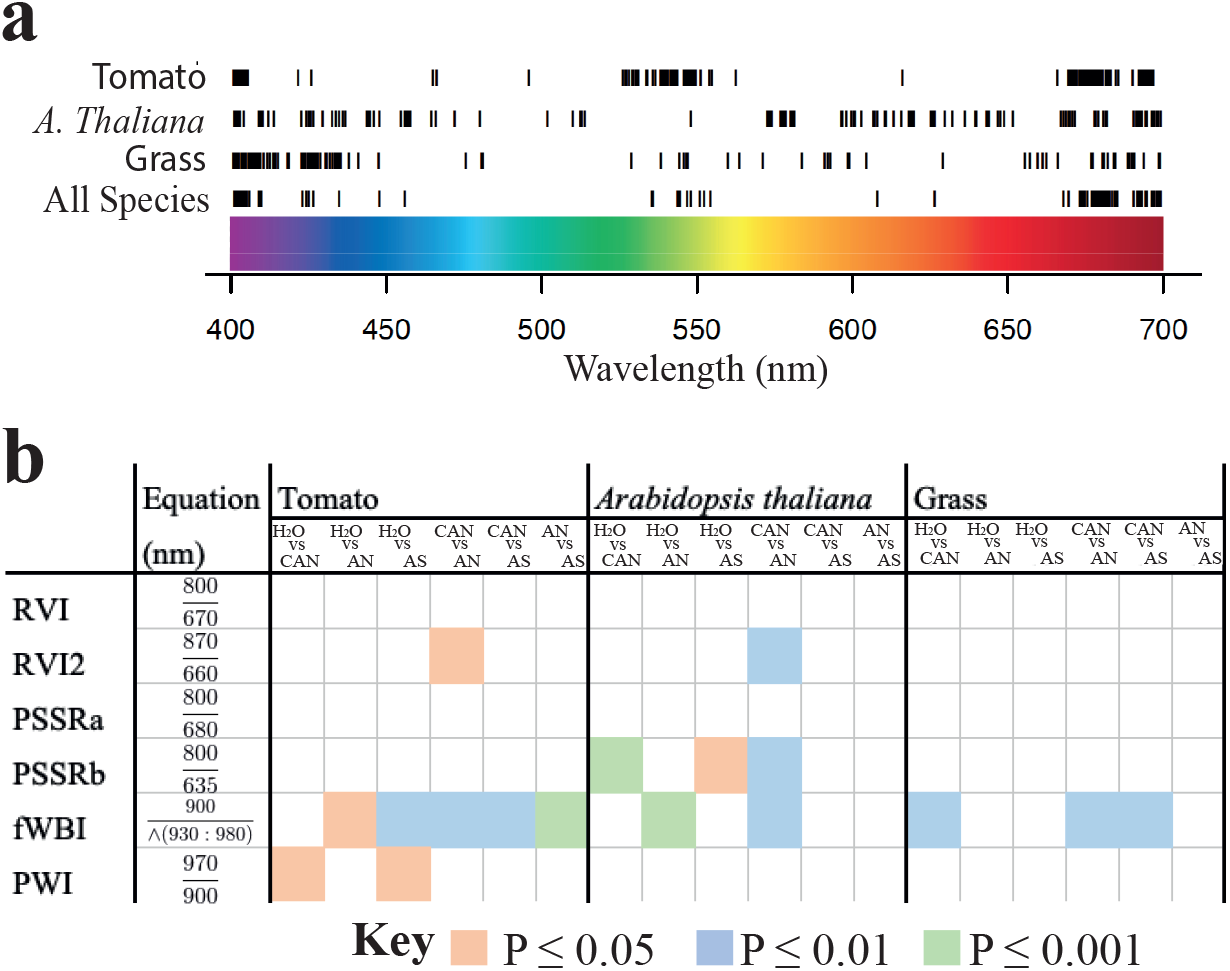
Ensemble regression identifies key spectral signatures that distinguish nitrogen treatments. **a** Ensemble regression of reflectance data from 400-700nm for tomato, Arabidopsis and lawn grass leaves 1-week after application of either 4mM of AN, CAN, AS or H_2_O, 1000 readings from 24 plants or grass swards per nitrogen treatment. Important wavelengths shown in black, selected for separate species if the absolute sum of regression coefficients is greater than two-standard deviations from the mean. The consensus all species bands were selected if the absolute sum of regression coefficients of all experimental replicates, for all species, was greater than two-standard deviations from the mean. **b** Vegetation ratio indices generated from hyperspectral data used to monitor nitrogen content (RVI, RVI2), water content (PWI, fWBI) and chlorophyll content (PSSRa, PSSRb). A nested ANOVA and Tukey HSD test was performed for each species with the experiment as the block to compare nitrogen treatments using specific indices. Significant differences are colour coded by p-value.

We expected total nitrogen abundance to vary across comparisons with the water control and were curious to see if variable nitrogen uptake could influence treatment classification. Thus, we chose to assess nitrogen abundance using a range of reflectance ratios that have previously been used to estimate nitrogen abundance in leaves such as RV1 and RV2 (D. M. El-Shikha *et al*., 2008; Zhu *et al*., 2008). Overall, we found that these simple indices could not be used to reliably distinguish any treatments groups (Fig 2b, Supplementary Fig 5).

However, reflectance ratios developed to estimate chlorophyll abundance via absorption of chlorophyll b (PSSRb-800nm/635nm) (Blackburn, 1998) could in some cases distinguish treatments, but not measures of chlorophyll a abundance (PSSRa-800nm/680nm) (Blackburn, 1998). Unusually, a water index measurement (fWBI-900nm/min(930:980nm) (Strachan, Pattey and Boisvert, 2002) could partially resolve some comparisons across our species, however this was not recapitulated with PWI (970nm/900nm) another water content estimator (Penuelas *et al*., 1997). This shows that our nitrogen treatments cannot be fully distinguished using these simplified ratio comparisons, while our regression approach and PCA analysis suggest multiple EM regions are needed to distinguish our nefarious nitrogen sources and controls.

### Customised multispectral imaging reliably distinguishes nitrogen treatments

PC and ensemble regression analysis suggest that numerous regions within the VIS-EM are needed to separate nitrogen treatments. So, we devised a highly customisable multispectral image analysis pipeline to acquire and analyse image data for plant leaves exposed to different nitrogen sources. We used simple colour pass filters that selectively transmit light within regions of interest identified by the ensemble regression analysis. Colour pass filters are placed in front of a smart phone camera during image acquisition (Supplemental Fig. 6). Images for multiple filters are registered and combined into a multichannel image stack. We then train random forest pixel and object classifiers (Sommer *et al*., 2011) on these stacks to build a machine learning system capable of distinguishing plants exposed to different nitrogen sources (Fig. 3 a, b & c). An initial pixel level classifier is first trained to distinguish plant leaves or leaf regions from background. This defines leaves as objects. An object classifier is then used to classify leaves by nitrogen treatment. This is trained using 2-6 regions of grass swards or tomato or Arabidopsis leaves from each treatment. This training data is then used to predict the treatment of the remaining leaves in the image.

**Figure 3.**
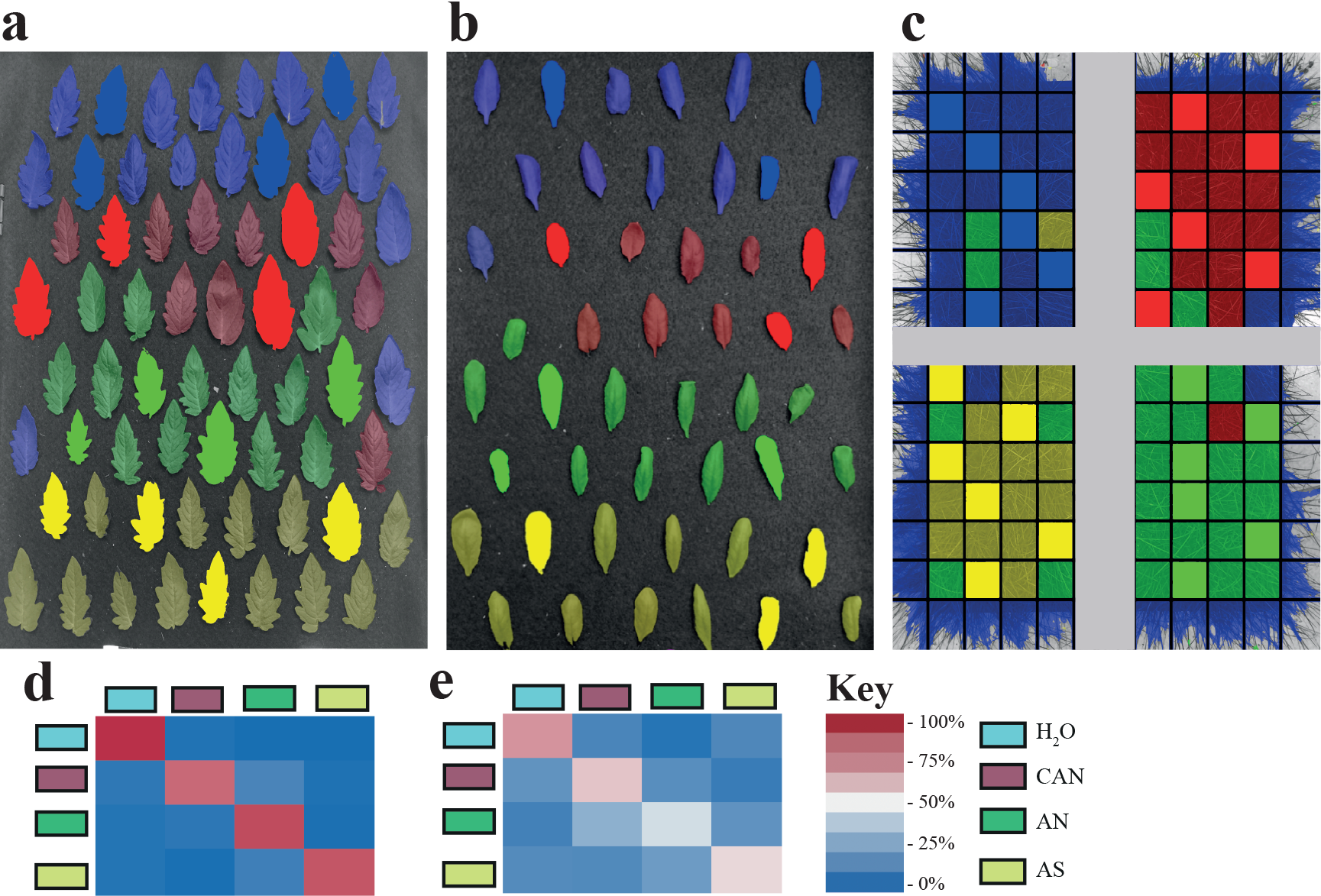
Machine learning classifiers built from multispectral data demonstrate superior performance. Multispectral image classification of **a** tomato, **b** Arabidopsis and **c** grass. Images acquired 1-week after application of either 4mM of AN, CAN, AS or H_2_O. Images were stacked and registered producing a multispectral image data cube. A random forest pixel classifier was used to remove background pixels and define leaf objects. A random forest object classifier was then used to predict nitrogen treatment in leaves. **a** 2-3 leaves, **b** 3-4 leaves and **c** 4-6 grass sward regions per treatment were used as training data. **a**&**b** rows 1&2 – H_2_O control; 3&4 - CAN; 5&6 - AN; and 7&8 - AS. **c**: top-left – H_2_O; top-right - CAN; bottom-right - AN; and bottomleft - AS. Predictions are blue-H_2_O control, red-CAN, green-AN, yellow-AS with training labels shown in solid colour. Confusion matrices showing mean accuracy for **d** multispectral and **e** RGB data for 3 experimental replicates from each species (9 experiments in total). Overall accuracy for multispectral classifier AN = 89%, AS = 86.51%, CAN = 81.18%, H_2_O = 94.79%. Accuracy of a machine learning classifier built using RBG images AN = 44.76%, AS = 54.28%, CAN = 58.36%, H_2_O = 70.12%. Accuracy was significantly different between RGB and multispectral images determined with a t-test (AN - P = 0.018; CAN - P = 0.049; AS - P = 0.00029 or H_2_O - P = 0.014).

The object classifier was capable of predicting nitrogen treatment with impressive accuracy (Fig 3d). Furthermore, multispectral image data created a classifier with significantly higher accuracy than models built using standard 3-channel RGB images, determined with a t-test (AN p = 0.018; CAN p = 0.049; AS p = 0.00029 or H_2_O p = 0.014). This shows that the regression analysis was useful in helping to target VIS-EM regions for subsequent multispectral analysis, with multispectral data providing superior classification performance compared to simple RGB images.

### Untargeted metabolomics identifies shifts in metabolites between nitrogen treatments

Our hyperspectrometry and multispectral image analysis show clear shifts in reflectance across the VIS-EM We reasoned this is likely due, in part, to differences in levels of chlorophyll and other nitrogen rich metabolites. However, we also reasoned that our treatments may alter plant secondary metabolism not directly influenced by nitrogen abundance. Irrespective of this we expect that shifts in chromophores across treatments influenced VIS-EM reflectance. To investigate this, we performed untargeted metabolomics on plant leaves one week after treatment with either 4mM AN, CAN, AS or H_2_O. Leaf tissue extracts were run on a HPLC-Q-TOF-MS according to De Vos *et al*. (De Vos *et al*., 2007) with the acetonitrile gradient modified to 100%. Monoisotopic masses of detected metabolites were filtered for count values >5000, fold change ≥ 2 and significant after a one-way ANOVA (P < 0.05) in all replicates per nitrogen treatment when compared to either Water or CAN as a control. PCA performed on differentially expressed monoisotopic masses (DEMMs) across biological replicates for each species show DEMMs group based on nitrogen treatment (Supplementary Fig 7). Hierarchal clustering was used to compare DEMMs. This compared expression for 163 DEMMs in Arabidopsis, 21 DEMMs in tomato and 31 DEMMs in grass with CAN as a control against AN and AS (Fig 4 a-c). Secondly, we compared expression for 72 DEMMs in Arabidopsis, 18 DEMMs in tomato and 62 DEMMs in grass for Water control against CAN, AN and AS (Fig 4 d-f). AN and AS appear more similar to each other compared to CAN and water control.

**Figure 4.**
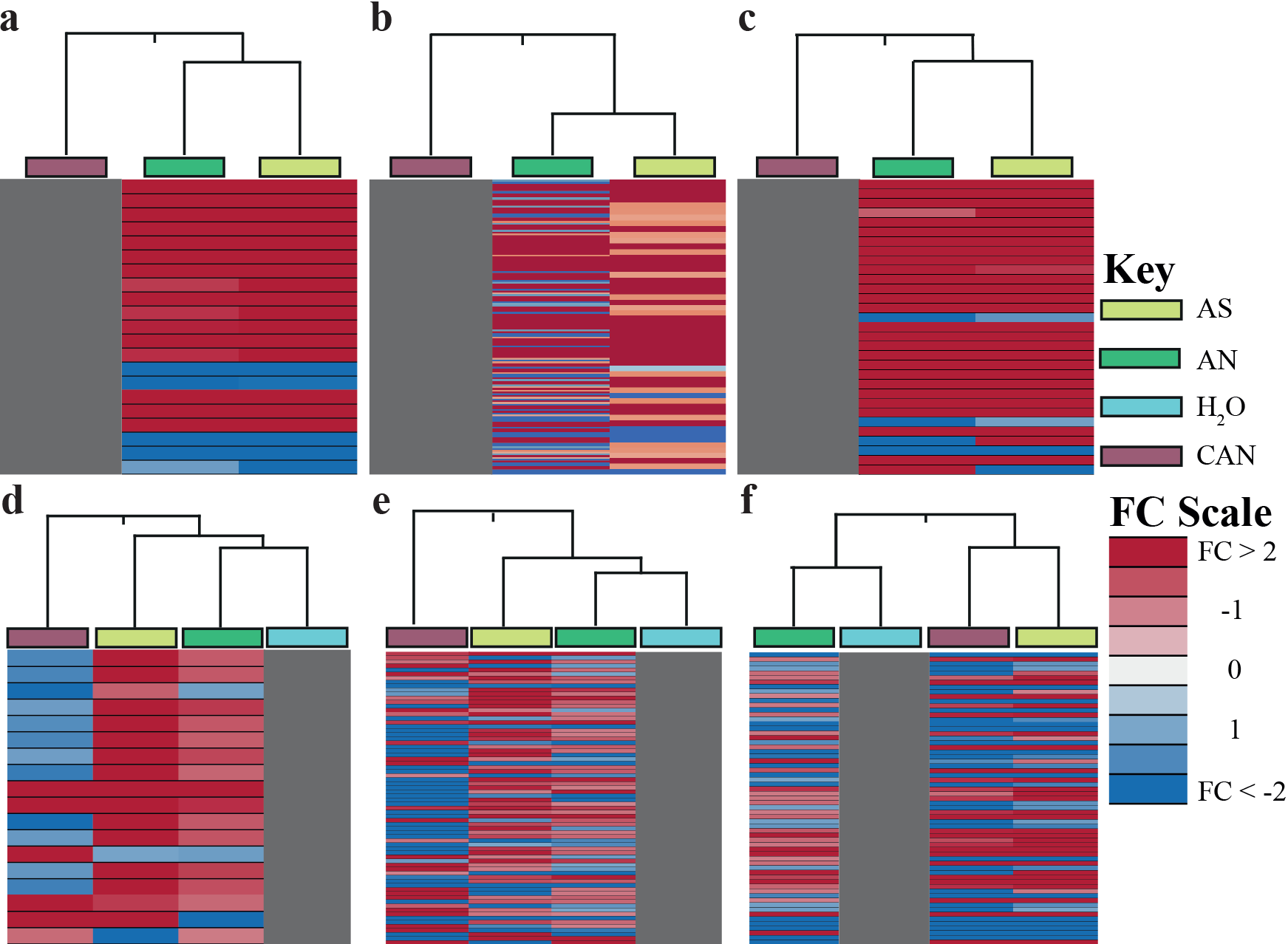
Hierarchical clustering of DEMMs reveals similarities between AS and AN treatment. tomato - **a** & **d** -tomato; **b** & **e** -Arabidopsis; and **c** & **f** - grass 1-week after application of either 4mM of AN, CAN, AS or H_2_O. 3 biological replicates per experiment and 2 experimental replicates per species. Extracted metabolites were run on HPLC-TOF-MS, Monoisotopic masses were filtered (count value >5000, fold change ≥ 2, significant after a one-way ANOVA p-value < 0.05) in all replicates per nitrogen source. **a**, **b** & **c** CAN or **d**, **e** & **f** H_2_O used as a control.

We searched for DEMMS in species specific PlantCyc databases (Schläpfer *et al*., 2017) using a neutral monoisotopic mass tolerance of ±50ppm. The majority of the DEMMs had no match (Fig. 5 a). However, we identified 21 differentially expressed known metabolites (DEKMs). Certain DEKMs classes contain chromophores known to influence plant colour, such as flavonoids (Panche, Diwan and Chandra, 2016) and light absorption, such as porphyrins (Lee *et al*., 2018) (Fig 5b). Porphyrins (a key component of chlorophyll (Mauzerall, 1977; Lee *et al*., 2018)) were upregulated in tomato leaves applied with AS compared to CAN. This supports our regression approach which identified the green (500-575 nm) region of the VIS-EM as key in helping distinguish variation between nitrogen treatments (Fig 2a).

**Figure 5.**
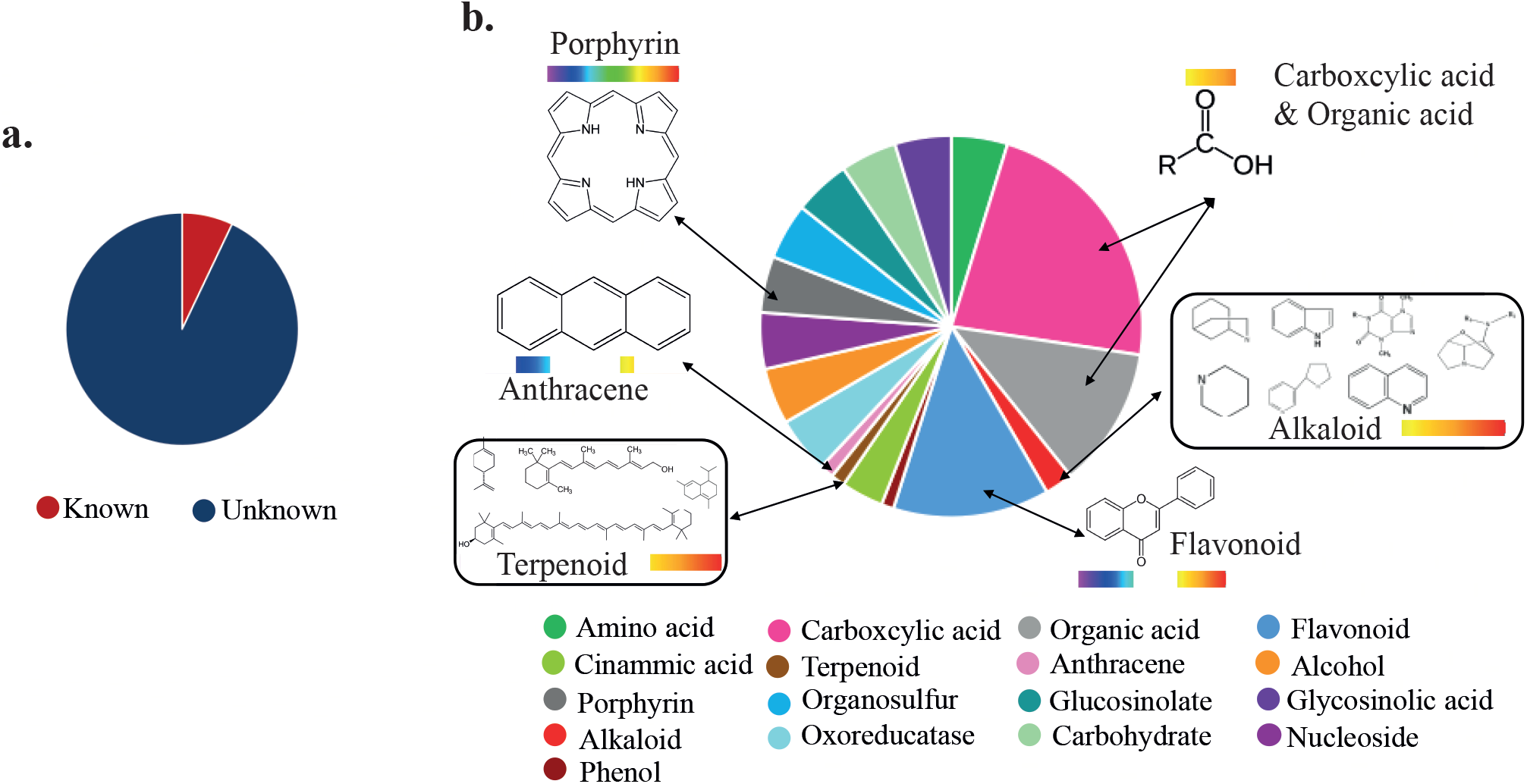
Differentially expressed metabolites include known chromophore containing molecules. DEMMs matched to Plantcyc(Schläpfer et al., 2017) using a mass tolerance of ±50ppm. **a** Most of the DEMMs identified across all species across all nitrogen treatments were unknown. **b** Pie chart of classes of the known metabolites. A total of 21 known metabolites were identified, each with a weighting of 1 in the pie chart. If a known mass could be more than a single metabolite group the value was divided accordingly among the possible groups. General structures shown for chemical classes containing known chromophores capable of influencing VIS-EM reflectance.

One specific DEKM is kaempferol-3-rhamnoside-7-rhamnoside a yellow flavonoid previously isolated from delphiniums (Kim *et al*., 2019). This DEKM is down regulated in Arabidopsis treated with AN and AS and up regulated with CAN compared to H_2_O. Yellow bands VIS-EM (560-590 nm) regions are highlighted by our ensemble regression analysis in Arabidopsis and lawn grass. Another DEKM, 3,8-divinyl protochlorophyllide is involved in chlorophyll and porphyrin biosynthesis (Nagata, Tanaka and Tanaka, 2007) and was upregulated in grass treated with AS vs H_2_O. Additionally, antrhone noranthrone was also upregulated in AS vs H_2_O, this DEKM is likely to be yellow as metabolites in the same structural family are yellow (Morse, 1947). Another yellow DEKM was the carboxcylic acid 1-O-sinapoyl-β-D-glucose, this DEKM was upregulated in CAN against H_2_O and downregulated in AS and AN compared to CAN. Interestingly, 1-O-sinapoyl-β-D-glucose is known to be involved in phenylpropanoid synthesis (Strack, 1982). Phenylpropanoids are the starting point for the biosynthesis of flavonols, isoflavanoids and anthocyanins (Nanda *et al*., 2017), all of which are known to regulate plant pigments across the spectra from purple and blue through to red and yellow (Forkmann and Martens, 2001; Katsumoto *et al*., 2007; Nakatsuka *et al*., 2007; Chiu *et al*., 2010; Falcone Ferreyra, Rius and Casati, 2012).

In conclusion, we have developed a sensitive multispectral image analysis machine learning classifier capable of distinguishing plants exposed to various nitrogen sources that are known to be involved in the production of illicit substances. We found that multiple regions within the VIS-EM are critical for facilitating nitrogen class distinction in a way that more simple reflectance ratios were incapable of doing. Exposure to different nitrogen sources induces differential expression of an array of metabolites including chromophore containing molecules. This helps to explain why the ensemble regression analysis identified numerous spectral regions across the VIS-EM as key for distinguishing nitrogen treatments.

We developed the system to facilitate detection and monitoring of illicit activities that utilise nefarious nitrogen sources, such as explosives and narcotic manufacture. Our results suggest significant potential for developing lab and remote sensing-based solutions to help extend this more fully. Also, this study highlights how plants interact differently with different nitrogen sources altering leaf biochemistry and suggests variable bioassimulation of nitrogen compounds. This could have important consequences in future studies of ecological and agricultural systems and biogeochemical cycling.

## Materials and Methods

### Plant Growth

Three different plant species were used in this study; *Arabidopsis thaliana* Col-0 (Arabidopsis), *Solanum lycopersicum* moneymaker, Suttons Seeds - tomato and mix of *Panicum* present UK lawn grass seed mix (Multipurpose lawn seed, Evergreen). Arabidopsis seeds where placed in 0.01% agar (Sigma Aldrich, UK) then at 4°C for 3 days before sowing, grass and tomato were sown from the packet. Seeds were planted in John Innes No.1, calcerite and perlite (6:1:1). Plants were grown under a 16-hour light dark cycle, with illumination of 150*μE m^-2^s^-1^* at 23°C and watered when needed. 1-week-old grass, 1.5-week-old Arabidopsis and 2-week-old tomato were watered with a single 4mM application of either calcium ammonium nitrate ((NO_3_)_3_CaNH_4_), ammonium sulphate ((NH_4_)_2_SO_4_), ammonium nitrate (NH4NO3) or water. One-week later leaves chosen at random or whole plants in each sample group were used for reflectance analysis or frozen in liquid nitrogen for metabolomic analysis.

### Reflectance spectrometry

Hyperspectral reflectance data was collected from 200 - 1025nm at a 0.4nm resolution, with a Flame-S-XR1, Ocean Optics, using a constant light source (DH-2000-BAL, Ocean Optics) and Ocean view software (Ocean Optics). The software was set to average 10 acquisitions for each spectrum recorded correcting for electric dark and non-linearity with a boxcar width of 2. 100 readings were taken per treatment group per experiment.

Hyperspectral data analysis was performed in R (Team, 2017), including and the adapted ensemble regression approach (Feilhauer, Asner and Martin, 2015). The ensemble regression was adapted to identify wavelength variance within and across species that significantly differed between nitrogen treatments. The reflectance intensity of the sample is first converted to a weighted coefficient before regression so a reflectance shift from 100 to 110 is weighted greater than a shift of 1000 to 1010, as the former is a 10% increase and the latter a 1% increase. Wavelengths were then selected for separate species if the absolute sum of regression coefficients is greater than two-standard deviations from the mean. The consensus all species bands were selected if the absolute sum of regression coefficients of all experimental replicates in the condition was greater than two-standard deviations from the mean.

### Multispectral image acquisition

Multispectral Images were taken in direct or simulated sunlight (iPhone 5s, Apple) using colour pass filters placed in front of the phone camera lens with no further modification. The colour filters used were #12, #23, #39, #65, #3407, #4990, #2005, #3441, #3202, #4290, #4690, #4460, #4790, #318, #376, #41, #378, #77, #93, #2002 (roscolux). Filters were chosen that selectively filtered across EM regions of interest identified in the regression analysis. Several filters cover redundant regions but also selectively filter within different EM regions. This results in an information rich data cube where single channels highlight intensity variance across some but not all treatment groups.

Multispectral image data cubes were constructed by registering images. Image registration was performed in Matlab. The images were separated into red, green or blue single-channel tif images. Tif images where then filtered by eye selecting only tif images with meaningful contrast to the background and/or between leaves across treatments. Selected tifs were stacked into a multispectral image stacks and processed in Ilastik (Sommer *et al*., 2011). A pixel level classifier was built to distinguish plant tissue from background. The background channel data was discarded, and the leaf channel data segmented into separate objects (σ = 1.2, making each leaf or grass sward a separate object. An object classifier was then used to classify leaves by nitrogen treatment which was trained using 2-6 regions of grass swards; tomato leaves; or Arabidopsis leaves from each treatment. This training data was then used to predict the treatment of the remaining leaves in the image using a random forest classifier using single and neighbourhood pixel (x = 40, y=40) intensities, defects and sharpness. This was used to generate predictions for each nitrogen treatment for the unlabelled image leaf objects.

### Metabolomics

1-week after the application of nitrogen treatments leaves were frozen in liquid nitrogen. Metabolites were extracted from either single frozen tomato leaves; 1g of pooled frozen grass leaves; or 1g of pooled frozen Arabidopsis. Metabolite extraction was performed according to De Vos *et. al* (De Vos *et al*., 2007) modified by extending the acetonitrile gradient to 100%. Extracted metabolites were run on HPLC-(129-, Agilent, Santa Clara, CA), Q-TOF-MS (6650, Agilent). Each nitrogen treatment had 3 biological replicates per experiment with 2 experimental replicates overall.

Metabolomics data was analysed using mass profiler professional (Agilent). Identified metabolites were filtered to have a count value greater than 5000 and a fold-change greater than 2 compared to the control; the control used was CAN against AN and AS or water as a control against CAN, AN and AS. Masses were then filtered to be significant after a one-way ANOVA (P < 0.05). Filtering was performed to ensure the differentially expressed metabolite was present in the above conditions in all 6 biological replicates from the 2 experimental replicates for the nitrogen treatment for the plant species. This was done for all three species separately.

Hierarchal clustering was performed on the significant differentially expressed monoisotopic masses in each plant species with Euclidian similarity distance and Ward’s linkage rule using CAN as a control against AN and AS or water as a control against CAN, AN and AS. Principal component analysis was performed on differentially expressed monoisotopic masses for each species. Differently expressed monoisotopic masses were used to query species specific databases (PlantCyc, mass tolerance = ±50ppm) (Schläpfer *et al*., 2017) to attempt to identify the metabolite.

## Supplementary information

**Supplementary figure 1.**
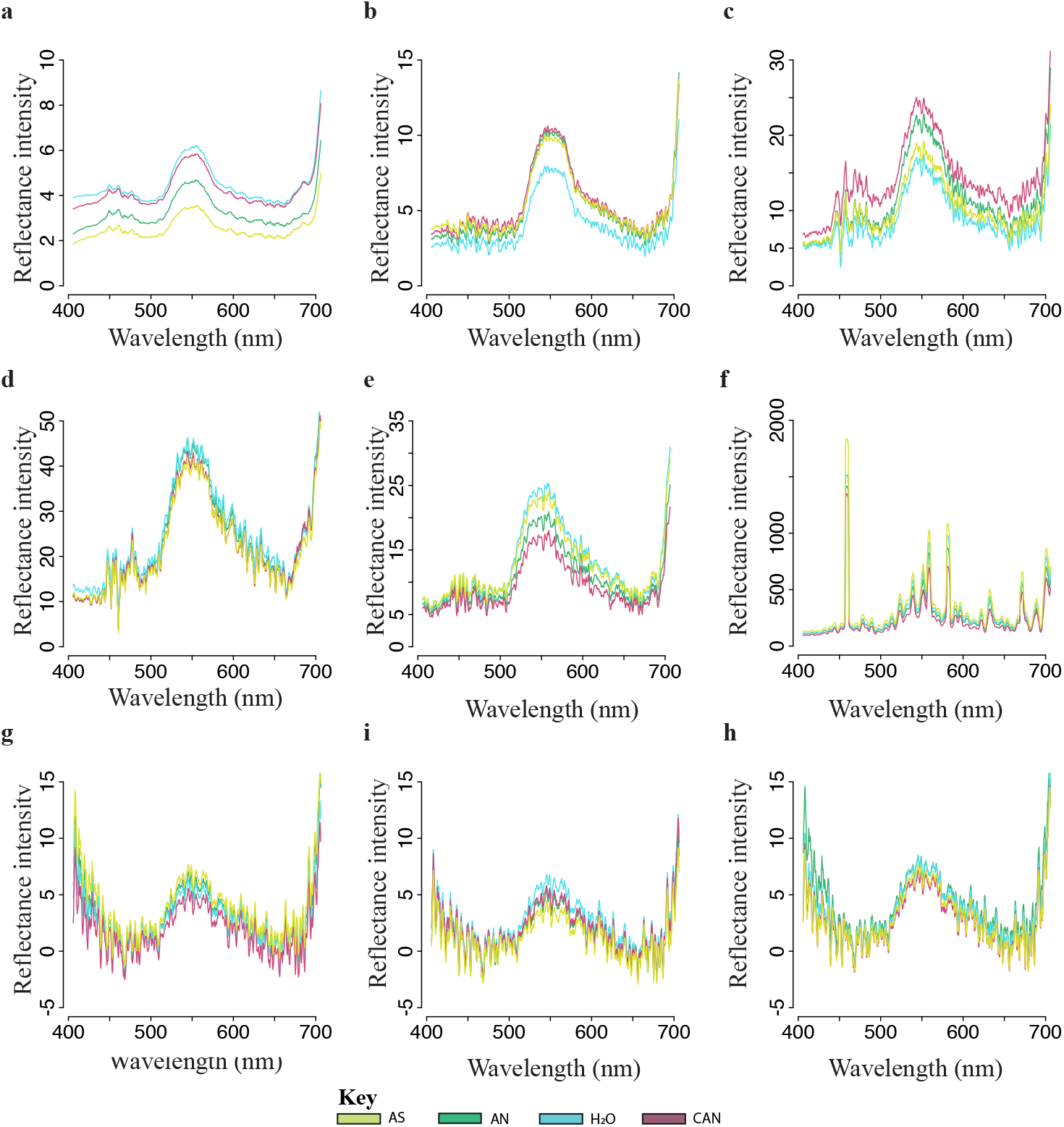
There is low visible hyperspectral reflectance pattern variance between across nitrogen sources. Mean reflectance intensity of the electromagnetic spectrum from 400-700nm for **a, b** & **c** tomato **d**, **e** & **f** Arabidopsis and **g**, **h** & **i** grass one week after application of either 4mM of AN (green), CAN (red), AS (yellow) or H_2_O (blue) (1000 readings, 24 plants or grass swards per nitrogen treatment, 0.4 nm resolution, boxcar width 2).

**Supplementary figure 2.**
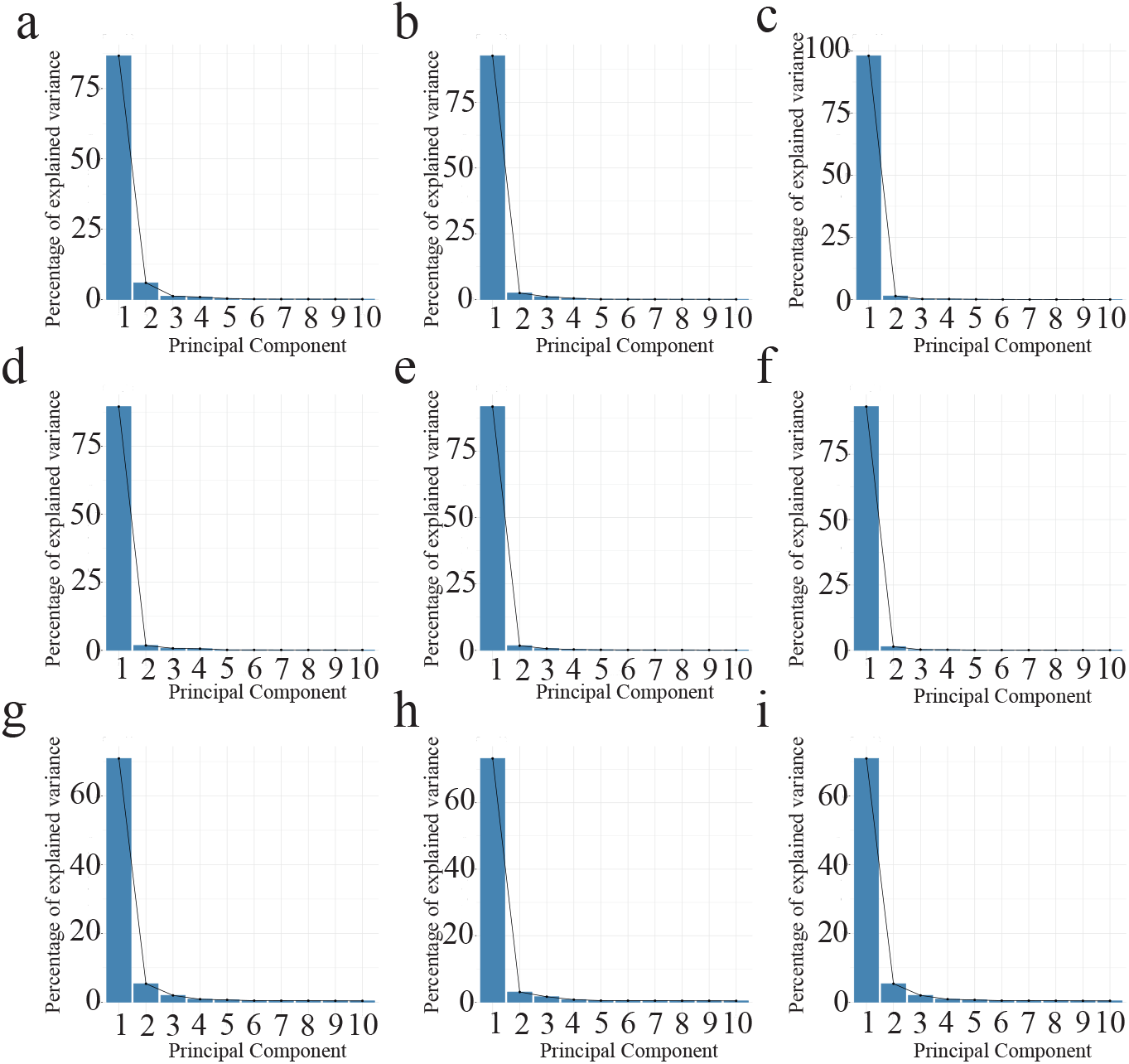
The majority of variance across data is captured in the first principal component. Principal component analysis was performed on the hyperspectral reflectance data from 400-700nm one week after application of either 4mM of AN (green), CAN (red), AS (yellow) or H_2_O (blue), 1000 readings taken from 24 plants per treatment. The percentage explained variance across all data captured by each principle component is shown for **a**, **b** & **c** Tomato; **d**, **e** & **f** Arabidopsis; and **g**, **h** & **i** grass. Three separate replicates shown for each species.

**Supplementary figure 3.**
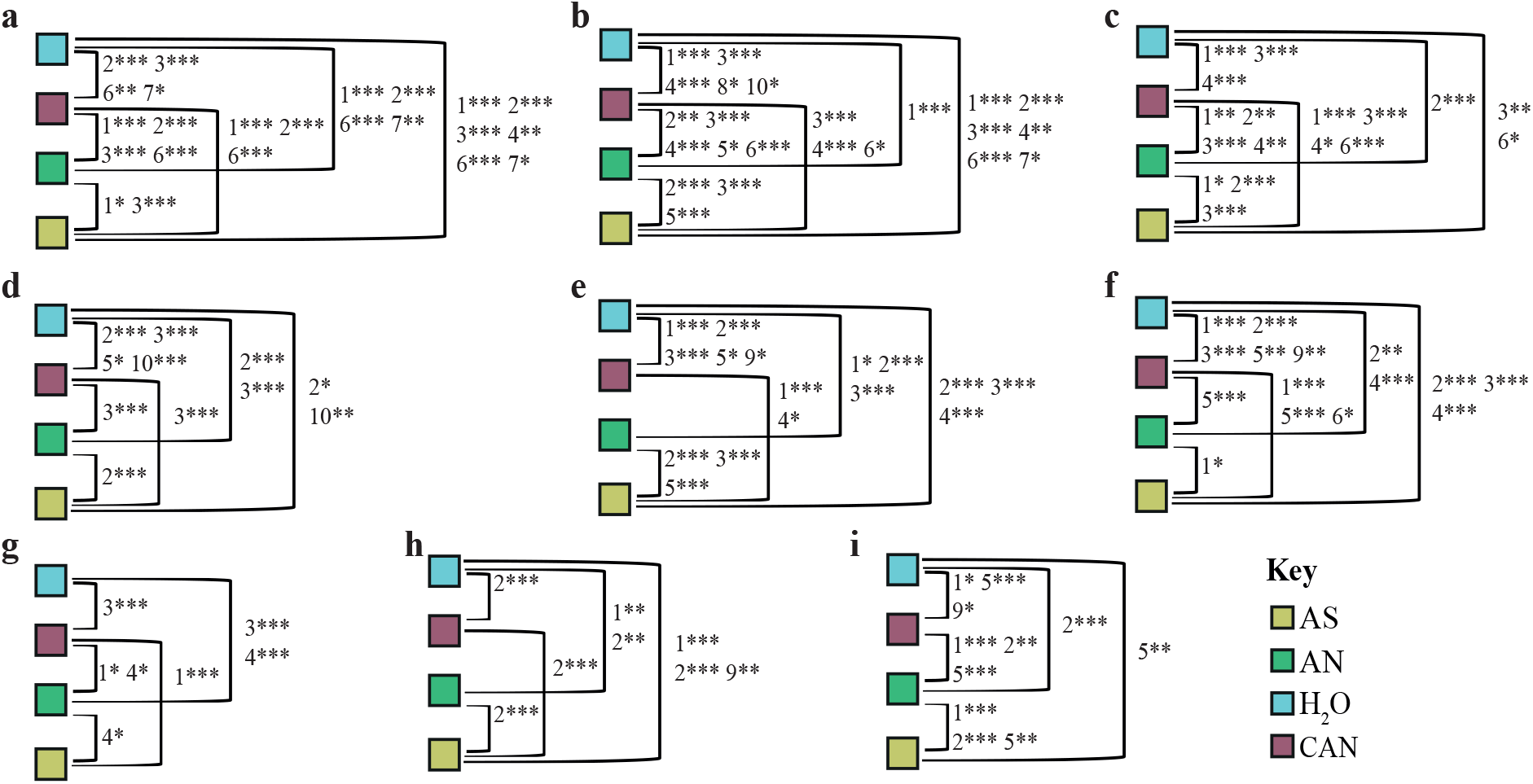
Hyperspectral variance reveals differences in plants treated with different nitrogen sources. Data for three replicates per species shown. ANOVA and Tukey’s HSD following PCA of reflectance data (400-700 nm) performed on PCs 1-10 for **a, b** & **c** tomato; **d**, **e** & **f** Arabidopsis; and **g**, **h** & **i** lawn grass one week after application of either 4mM of AN (green), CAN (red), AS (yellow) or H_2_O (blue), 1000 readings taken from 24 plants per treatment. ANOVA and Tukey’s HSD following PCA of reflectance data (400-700 nm) performed on PCs 1-10. Lines between groups are labelled with PCs which contain variation that can significantly distinguish treatment group pairs. * indicates p-value for Tukey’s HSD < 0.05, ** < 0.01 and *** < 0.001. With the exception of e, g & h ANOVA followed by Tukey’s HSD was able to resolve all nitrogen treatment comparisons across all replicates and species.

**Supplementary figure 4.**
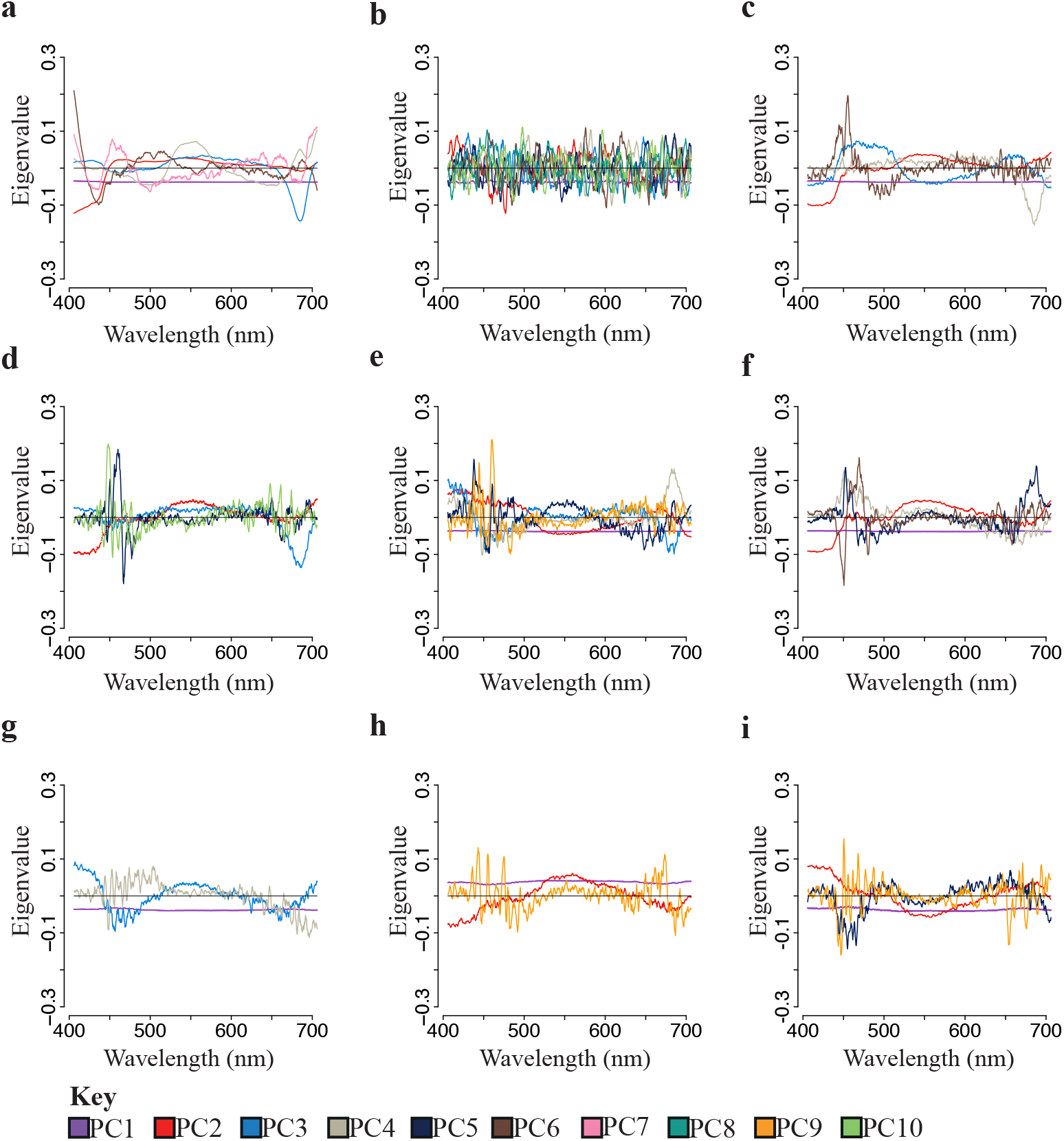
Loadings of principle components that could significantly separate nitrogen sources capture variation from regions identified by machine learning regression. Principal component analysis was performed on the hyperspectral reflectance data from 400-700nm one week after application of either 4mM of AN (green), CAN (red), AS (yellow) or H_2_O (blue), 1000 readings taken from 24 plants per treatment. **a, b** & **c** tomato; **d**, **e** & **f** Arabidopsis; and **g**, **h** & **i** lawn grass eigenvalue plots of PCs which could significantly explain variation between nitrogen treatments in hyperspectral data (400-700nm) after ANOVA and Tukey HSD test on single components (p < 0.05).

**Supplementary figure 5.**
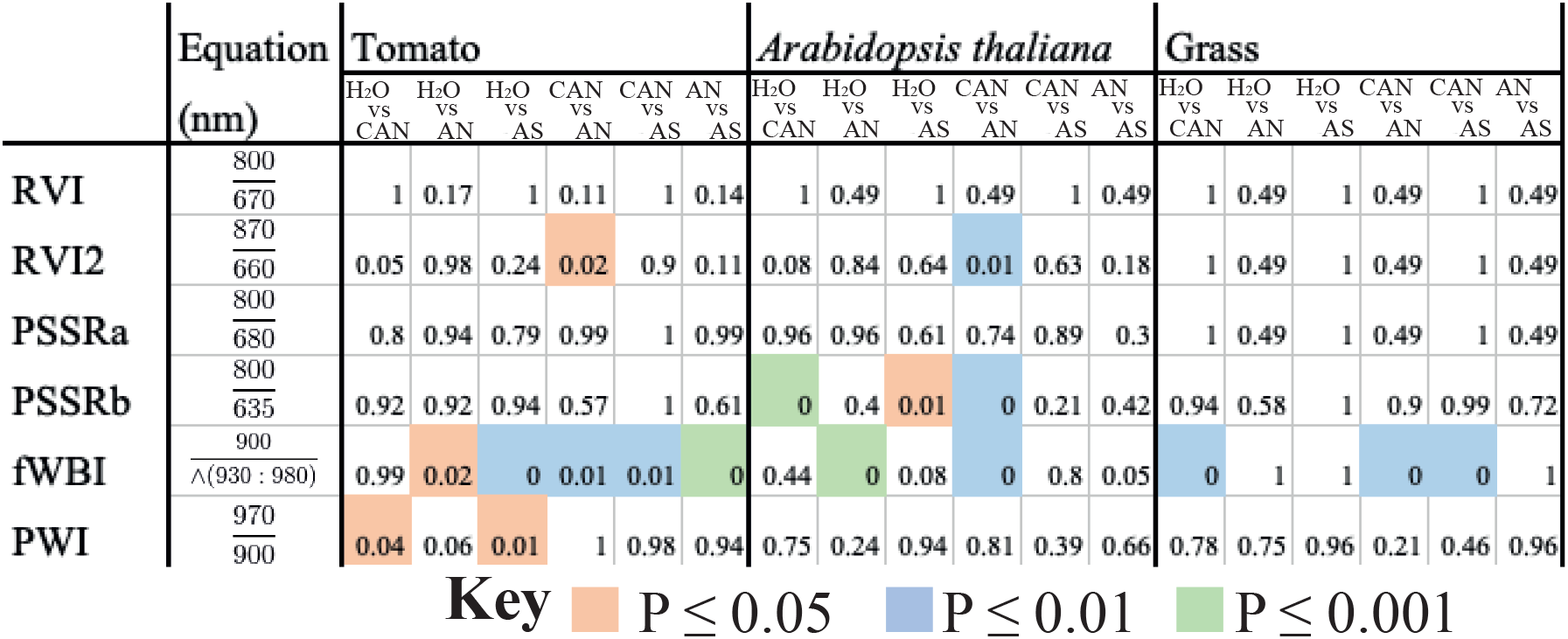
Plant reflectance indices are unable to consistently differentiate nitrogen treatments. Hyperspectral reflectance intensity was taken from either tomato, Arabidopsis and lawn grass leaves 1-week after application of either 4mM of AN, CAN, AS or H_2_O, 100 readings from 24 plants or grass swards per nitrogen treatment, for 3 experimental replicates. Vegetation ratio indexes that monitor nitrogen content (RVI, RVI2), water content (PWI, fWBI) and chlorophyll content (PSSRa, PSSRb) were then calculated for each experiment. This produced 100 index values per treatment per experiment. Then a nested ANOVA and Tukey HSD test (P values shown) was performed on the index values for each species with the experiment used as the block.

**Supplementary figure 6.**
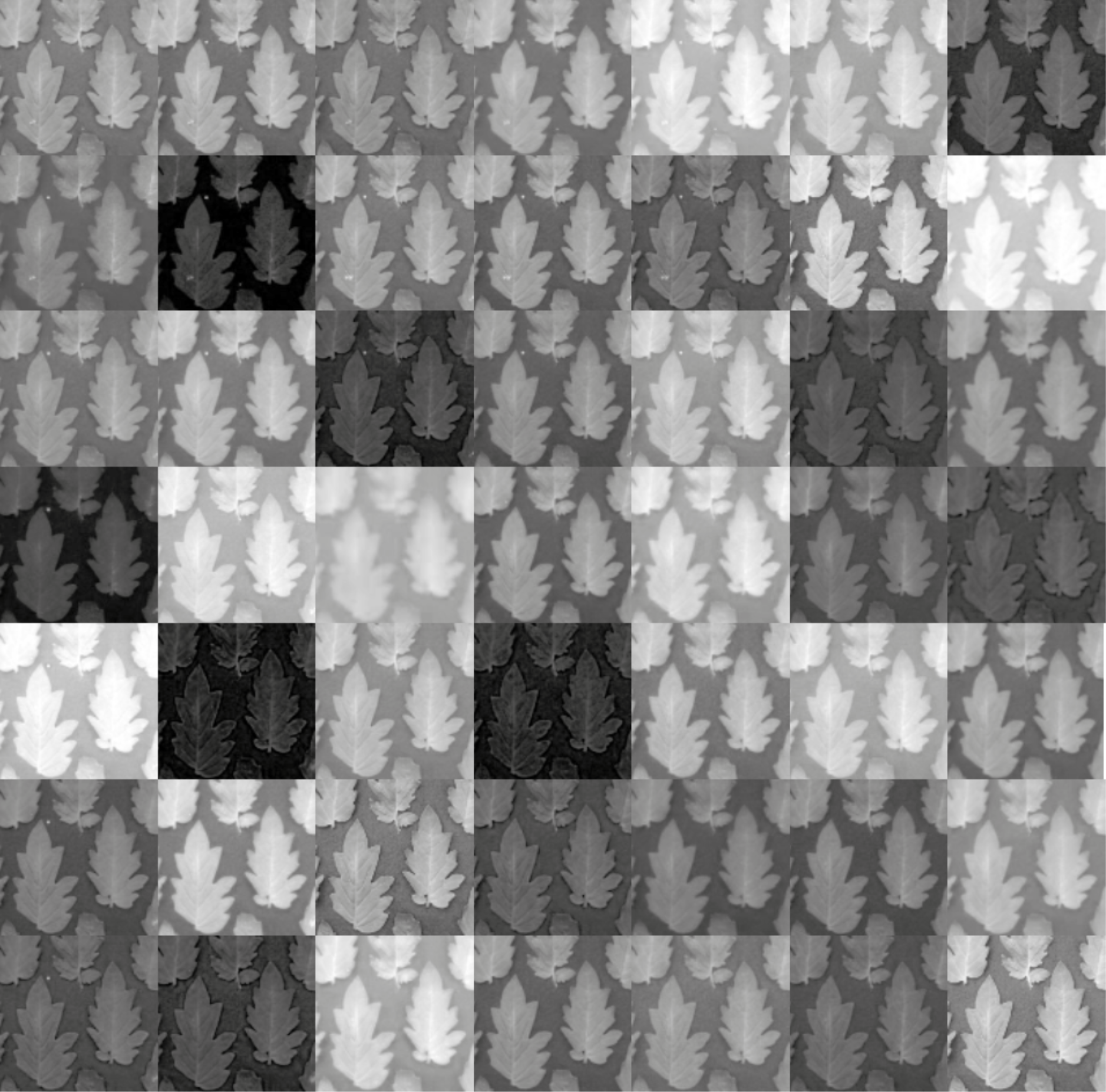
Cropped images of leaves captured using different filters used to form a multispectral cube for analysis. Cropped images of randomly selected tomato leaves from 1 experiment taken 1 week after the application 4mM of NH4NO with colour pass filters that selectively transmit light in regions of interest. This illustrates the intensity diversity across filters channels in the multispectral cube. Images were registered to align and then split into individual colour pixel channels. The pixel channels without contrast were discarded. The grid of images was then be stacked into a multispectral cube for analysis.

**Supplementary figure 7.**
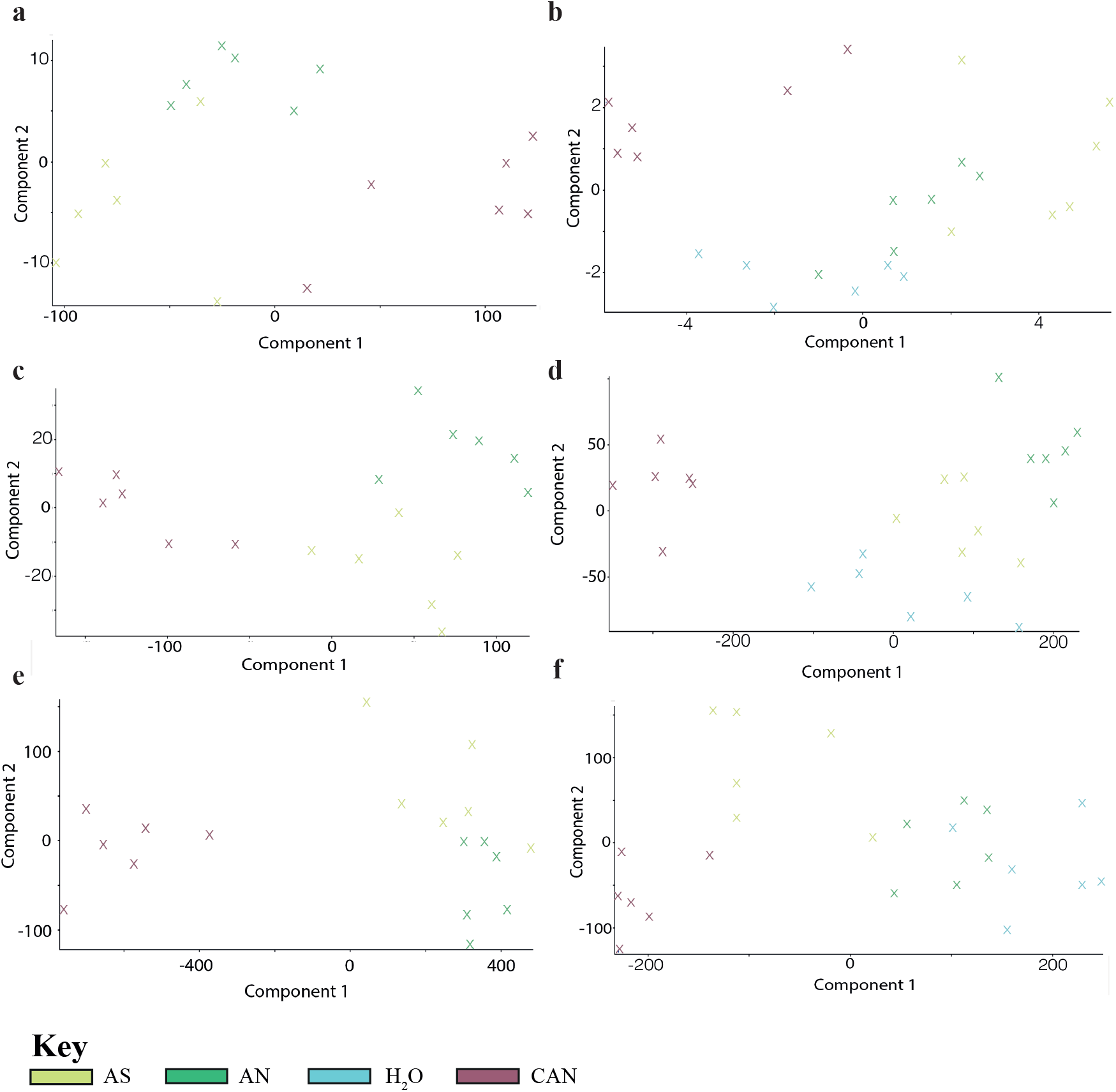
Significant monoisotopic mass expression identfied with untargeted metabolomics cluster after PCA dependant on nitrogen source applied to the plant. Untargeted metabolomics was performed on plant leaves one week after treatment with either 4mM AN, CAN, AS or H_2_O according to De Vos et al., (De Vos et al., 2007) with the acetonitrile gradient modified to 100%. Leaf tissue extracts were run on a HPLC-TOF-MS. Monoisotopic masses of detected metabolites were filtered for count values >5000, fold change ≥ 2 and significant after a one-way ANOVA (P < 0.05) in all replicates per nitrogen treatment when compared to the control. The control was CAN for AN and AS or water as a control against CAN, AN and AS. PCA was performed on differentially expressed monoisotopic masses in each experiment. The PC scores for the first component were plotted against the second for each experiment, for **a** & **b** tomato; **c** & **d** Arabidopsis; and **e** & **f** grass. The replicates visibly clustered based on the nitrogen treatment applied. Two experimental replicates performed per species.

## Author contributions

C.A. performed hyperspectral and multispectral data collection and data analysis, performed metabolite extractions and analyzed metabolomic data, helped design experiments, conceive the project and write the manuscript. S.C and D.B. performed the HPLC helped and advised with metabolomic data analysis. O.W. conceived the project, helped design experiments and wrote the paper.

## Acknowledgements

This work is was supported by the following grants: BB/M011178/1 and NE/M018768/1.

